# Intrinsic Degradation of the Type-II Antitoxin ParD from *Pseudomonas aeruginosa*

**DOI:** 10.1101/2021.03.29.437564

**Authors:** Kevin J. Snead, Christina R. Bourne

**Affiliations:** Department of Chemistry and Biochemistry, University of Oklahoma, Norman OK USA

## Abstract

Type-II Toxin Antitoxin (TA) systems are regulated by differential half-lives of the resulting non-secreted proteins, such that the neutralizing antitoxin undergoes continual degradation and replenishment to maintain neutralization of its cognate toxin. Antitoxin proteins are widely reported as labile, including upon purification and *in vitro* storage. During the course of studies on a ParDE TA system we noted a prevalent *in vitro* degradation of the ParD antitoxin. In efforts to combat this for practical use in assays, we characterized parameters impacting the degradation as well as the resulting products. These revealed a mechanism likely mediated by a serine or metal-dependent protease. Using Direct Infusion Mass Spectrometry, the cleavage products were identified as an essentially intact DNA binding region of the antitoxin and with the toxin binding domain completely removed. No other species were identified in the solution, such as a contaminant that may mediate such cleavage. Therefore, while our studies revealed viable strategies to mitigate the *in vitro* degradation they did not identify any protease, leaving open the possibility of a potential auto-catalytic proteolytic activity of the antitoxin proteins.

## Introduction

Type-II toxin antitoxin (TA) systems are composed of two proteins that impact of bacterial growth. The toxin is a non-secreted protein that inhibits essential cellular processes, resulting in disruption of normal cell growth. During normal growth conditions, the toxin is neutralized by the formation of a protein-protein complex with a cognate antitoxin protein, which sequesters the toxin from its cellular target (1,2). TA systems were first identified on plasmids, and were shown to control plasmid inheritance through post-segregational killing of daughter cells lacking the plasmid harboring the TA system (3-5). Post-segregational killing of plasmid free daughter cells is based on the susceptibility of antitoxins to degradation, resulting in shorter half-lives relative to their cognate toxins, allowing for accumulation of excess free toxin over time (1). The AAA+ proteases Lon and Clp have been shown to be involved in antitoxin turnover *in vivo* (6-8).

The antitoxin ParD belongs to the MetJ/Arc transcriptional repressor family (Pfam clan CL0057), and is composed of an N-terminal ribbon-helix-helix (RHH) DNA-binding motif and a C-terminal toxin-binding domain consisting of two alpha-helices (helix 2 and 3) which wrap around the ParE toxin resulting in an extensive protein-protein interaction (9-14). ParD antitoxins were originally shown to be disordered in the absence toxin binding, resulting in an unstructured toxin binding region, as shown in the NMR structure of the ParD antitoxin from the plasmid RK2 (10). In contrast, a chromosomal ParD from *Caulobacter crescentus* was shown by circular dichroism to be well-ordered in the absence of its cognate ParE (9). Secondary structure analysis by Fourier transform infrared spectroscopy of a chromosomal ParD from *Pseudomonas aeruginosa* indicates the C-terminal alpha-helix (helix 3) is unstructured in the absence of ParE binding (*unpublished data*). In addition to its intrinsic disorder, the RK2 ParD was shown to undergo *in vitro* degradation upon storage, resulting in highly specific degradation products (15). It remains unclear as to whether the degradation of RK2 ParD was caused by contamination of co-purifying cellular proteases, or an auto-catalytic function of ParD itself (15).

The current study examined a similar *in vitro* degradation upon storage of the antitoxin ParD (PA0125) from *Pseudomonas aeruginosa* with the aim of identifying the role of antitoxin stability in TA system regulation. During these studies we identified that *Pseudomonas aeruginosa* ParD undergoes highly specific degradation, similar to previous observations of the RK2 plasmid-derived ParD antitoxin (15). Assays were carried out in the presence of different protease inhibiting reagents, which revealed a moderate impact of an irreversible serine protease inhibitor (PMSF) or in the presence of EDTA. Inclusion of a “complete cocktail” of protease inhibitors, supplemented with EDTA was able to totally block the observed degradation. Based on the cocktail components, and the partial blockage with PMSF or EDTA, it appears the proteolytic cleavage may be mediated by either a serine or metal-dependent protease. Using direct infusion mass spectrometry, we mapped the specific site of antitoxin cleavage and determined that the ParD toxin-binding region is degraded in the absence of ParE toxin, leaving behind the intact DNA-binding domain of ParD.

## Results

### *Pseudomonas aeruginosa* ParD is unstable and prone to *in vitro*

Antitoxins have been reported as disordered or unfolded at the toxin-binding region and, as such, are prone to degradation in the cell; however, there are also observations of an inherent instability of the antitoxin protein even in a purified form (10,16,17). Throughout the course of these experiments, a marked instability of the purified wild-type ParD antitoxin *in vitro* was noted. To better understand the kinetics of this instability, wild-type ParD was assessed for degradation as a function of storage time by Tris-Tricine PAGE analysis (Fig. 1). Monitoring wild-type ParD over a 10-day timeframe reveals that ParD is intrinsically unstable and degrades over time. The rate of degradation is modestly affected by temperature, resulting in a shortened half-life of approximately 2.5 days at both 23°C and 37°C relative to ParD samples stored at 4°C with a half-life of approximately 6 days (Fig. 1).

**Figure 1.**
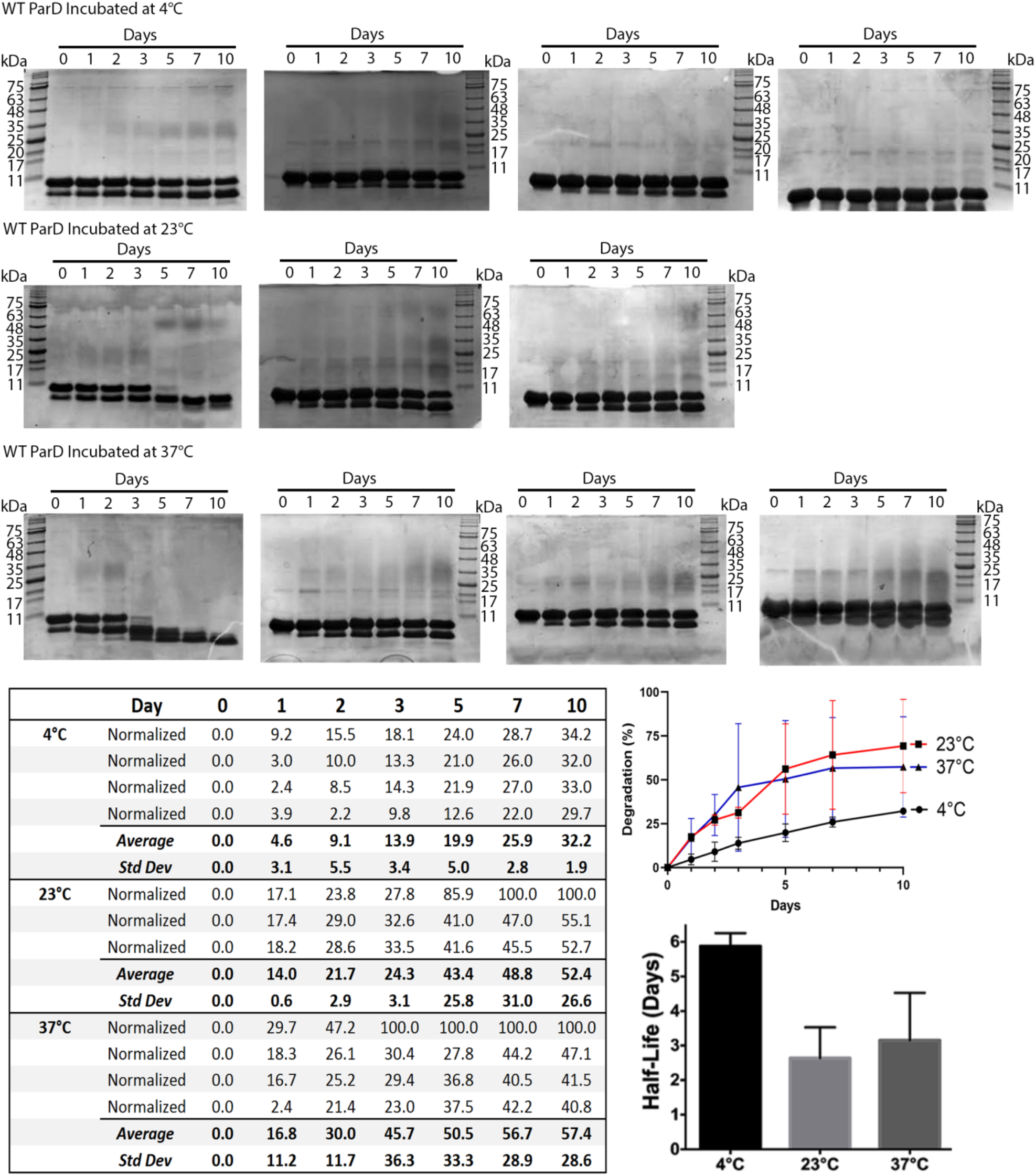
ParD degradation shows temperature and time dependence. *In vitro* degradation of ParD was monitored during storage at the indicated temperatures. Band intensities were measured using ImageJ, and degradation was calculated as a percentage of total intensity for each lane, and normalized to account for degradation present in the initial “Day 0” sample. A plot of the resulting values is also given.

### Degradation of ParD is blocked by inclusion of EDTA and protease inhibitor cocktail

To gain insight into the catalytic mechanism(s) of ParD degradation, we undertook a series of experiments to characterize potential inhibitors of the observed protein degradation when stored in a 50 mM Tris pH 8.5, 300 mM NaCl buffer. To ensure the observed degradation was not the result of contamination by fungal or bacterial cells, purified ParD was treated with 0.02% (w/v) sodium azide (NaN_3_). Addition of sodium azide had a small impact (∼ 28% reduction relative to the control sample) when assessed ten days post-purification (Fig. 2). While sodium azide had a small impact, the concentration of sodium azide used is sufficient for preventing both fungal and bacterial growth; indicating the degradation is unlikely to be caused contamination. The metal-chelating compound EDTA was also tested to potentially block the activity of any metal-dependent proteases. The addition of EDTA caused a significant decrease (∼ 50% reduction relative to the control sample) in ParD degradation when assessed ten days post-purification (Fig. 2). The irreversible serine protease inhibitor partially blocked degradation (∼ 40% reduction relative to control sample), while a commercially available protease inhibitor cocktail containing AEBSF, Bestatin, E-64, Pepstatin A, Phosphoramidon, Leupeptin, Aprotinin, and supplemented with EDTA blocked all degradation, showing no increased degradation after the initial “Day 0” time point (Fig. 2).

**Figure 2.**
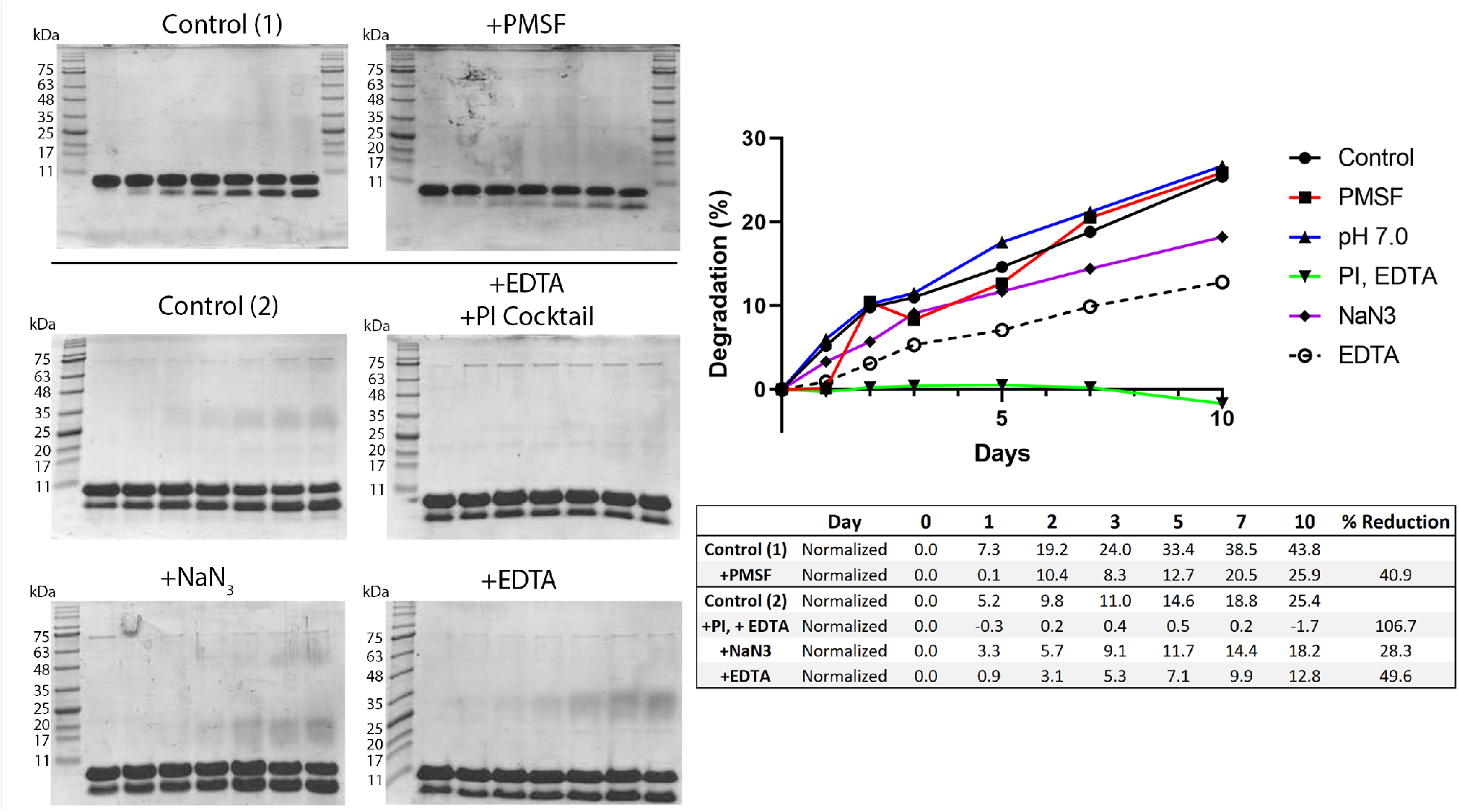
ParD degradation is blocked by a protease inhibitor (PI) cocktail. *In vitro* degradation of WT ParD was monitored during storage at 4°C, in the presence of inhibitors of protein degradation. Band intensities for ParD were calculated as a percentage of total intensity for each lane, and normalized to account for degradation present in the initial “Day 0” sample. A plot of the resulting values is also given.

### The toxin-binding region of wild-type ParD is degraded in the absence of ParE

Direct-infusion mass spectrometry (DIMS) was utilized to identify the remaining wild-type ParD fragment after degradation. Both degraded and non-degraded samples were directly injected into the ionization source and analyzed by mass spectrometry. Non-degraded wild-type ParD contained two prominent molecular weight species (10731.5 Da and 21462.98 Da), corresponding to ParD monomer and dimer respectively (Fig. 3). In comparing the mass spectrum of intact ParD to the resulting spectra of degraded ParD (over the same m/z window), multiple molecular weight species were evident and correlate to cleavage occurring in the region corresponding to ParD residues 42-47 (^42^RQQWQV^75^) (Fig. 4), which are located immediately after the ParD dimerization region. The most prevalent species (100% relative abundance) observed is a 5809.5 Da fragment resulting from cleavage at glutamine 46 of ParD (Fig. 6). Two additional high abundant species (60% relative abundance) were also observed with molecular weights of 5908.5 Da and 5266.4 Da (Fig. 4). The 5908.5 Da fragment results from cleavage at valine 47 of ParD. The 5266.4 Da fragment and an additional 5365.0 Da fragment (30% relative abundance) both result from cleavage at glutamine 43, with the smaller fragment resulting from the loss of the neighboring arginine 42 side chain (Fig. 4). These fragments indicate a relatively specific degradation pattern and implies that it could take place in a sequential manner. Further, it appears that the toxin-binding region of ParD is completely degraded, as no peaks corresponding to amino acids after residue 47 were evident, leaving only the ribbon-helix-helix (RHH) DNA binding domain of the antitoxin. These results are consistent with the gel analysis (Figs. 1-2), which shows accumulation of a single band of degraded protein, with no other detectable ParD fragments.

**Figure 3.**
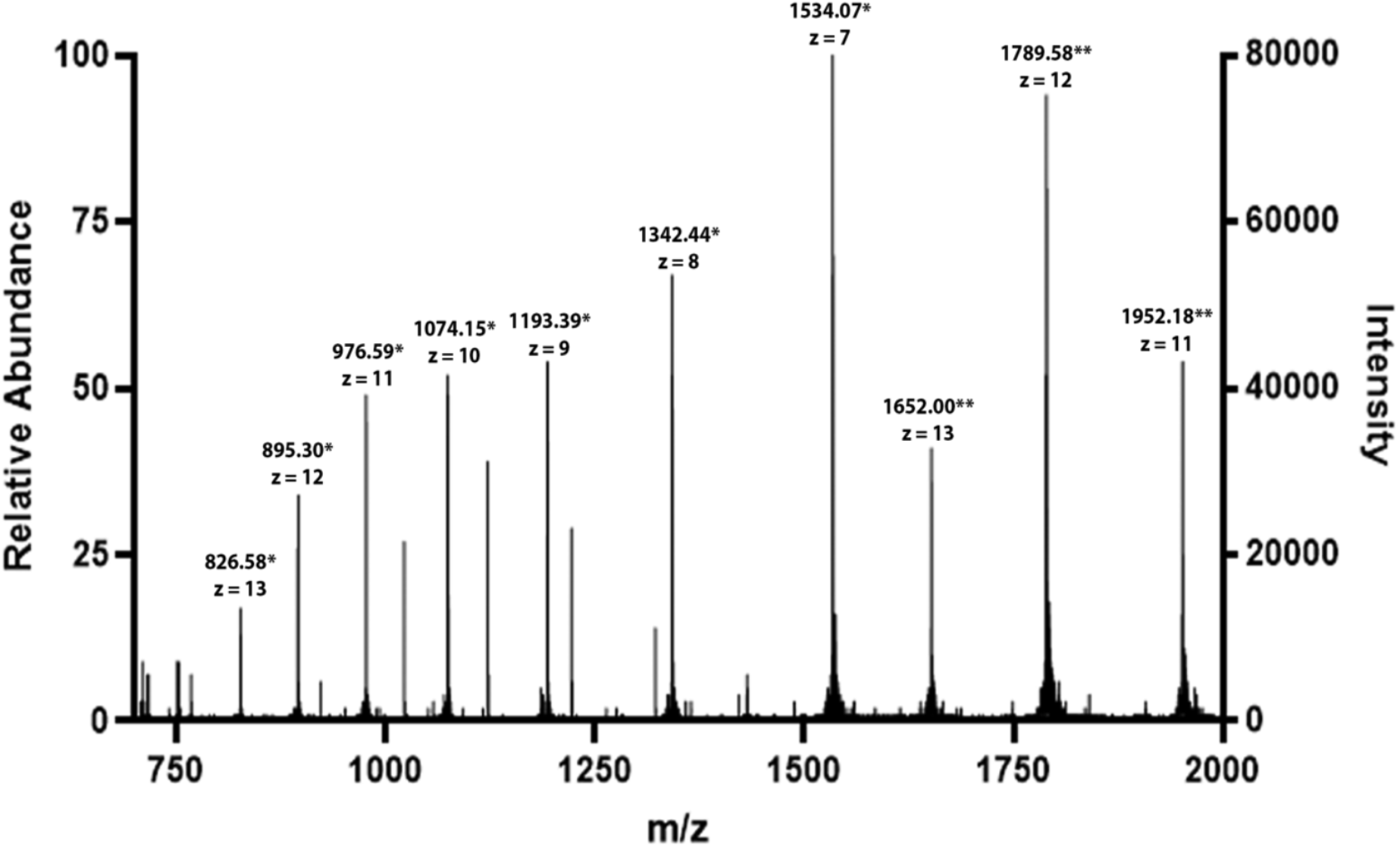
Direct infusion mass-spectrometric analysis of wild-type ParD before degradation reveals the major species are monomer and dimer. *Indicates monomeric ParD; **Indicates dimeric ParD. No additional species are evident within this mass window and ionization mode.

**Figure 4.**
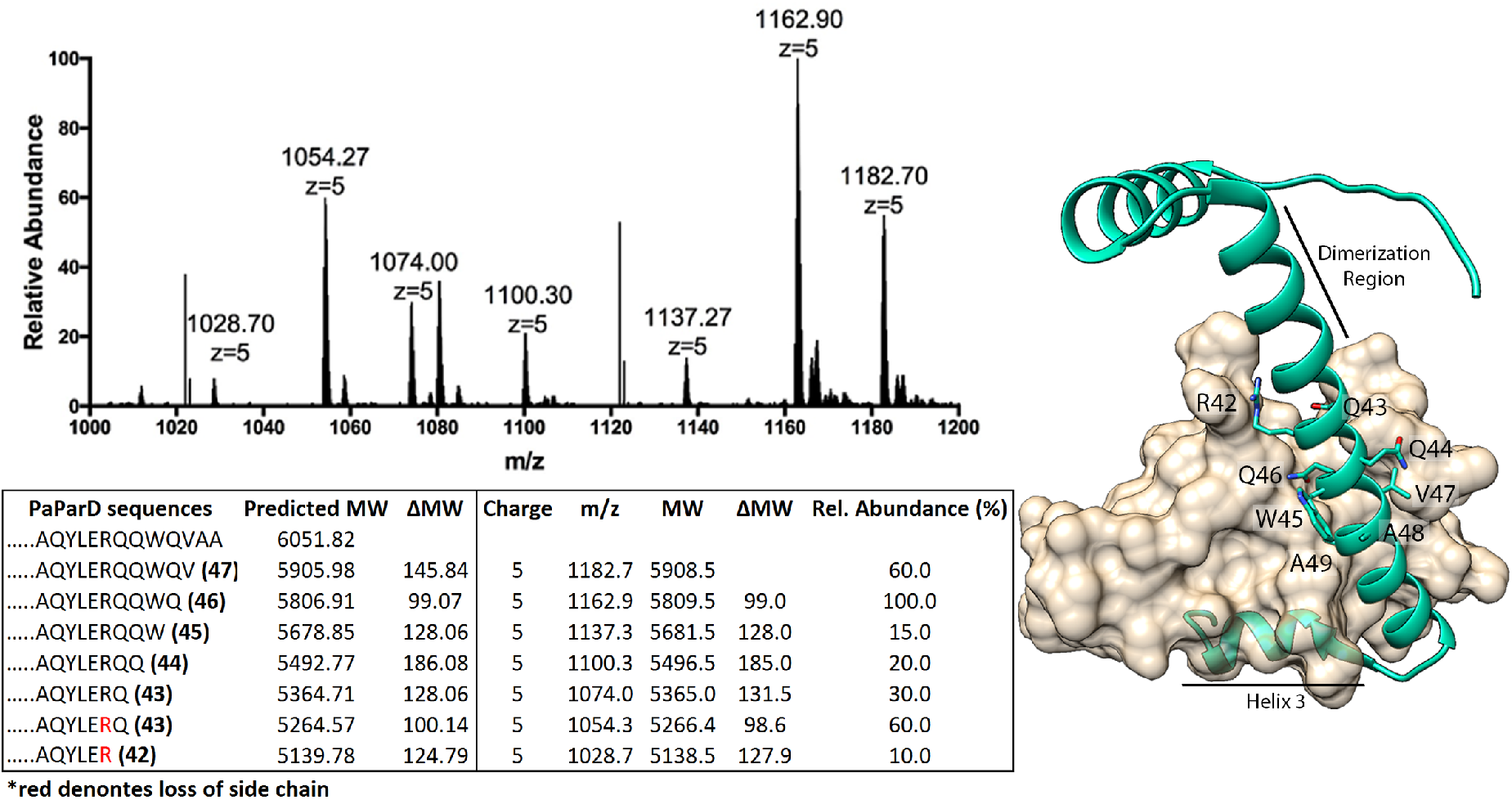
Intrinsic degradation of WT ParD results in complete loss of the toxin-binding region. Direct Infusion Mass Spectrometry of degraded ParD, +5 charged species shown (see **Fig. 3** for intact WT ParD mass spectrum). (Right) ParD (green) with amino acids corresponding to fragments detected by mass spectrometry displayed on the ParDE complex; ParE toxin is shown as a tan surface (PDB ID: 6XRW).

## Discussion

Inherent instability of antitoxins *in vivo* is a key rationale for the differing half-lives between toxins and antitoxins; however, the explanation for this instability relies on cellular proteases that are presumed absent in our *in vitro* storage experiments (10,16-18).Wild-type ParD from *Pseudomonas aeruginosa* shows discrete degradation products visible only a few days after purification (Figs. 1-2). Similar observations of *in vitro* degradation have been reported for the ParD antitoxin from the RK2 plasmid, where three prominent masses were identified when analyzed by MALDI-TOF mass spectrometry (15). Sequence alignment of RK2 ParD and *Pseudomonas aeruginosa* ParD indicates RK2 ParD contains a homologous sequence (^43^DQAWQ^47^) to the cleavage sites we have identified in *Pseudomonas aeruginosa* ParD (^42^RQQWQV^47^). The masses of the degradation products seen in RK2 ParD however, do not correspond to cleavage at this site. Instead, analysis of the fragments of RK2 suggests two distinct cleavage events resulting in a 6006 Da fragment corresponding to cleavage at leucine 54, and a 6873 Da fragment corresponding to cleavage at leucine 61. Since both RK2 ParD and *Pseudomonas aeruginosa* ParD have intrinsically disordered C-termini in the absence of ParE binding, the intrinsic degradation observed may be expected for all unstructured ParD proteins (10,15). The degradation of ParD can be effectively blocked by inclusion of EDTA or a protease inhibitor cocktail that contains inhibitors of serine, cysteine, aspartic, and metalloproteases. The partial inhibition of degradation by both the irreversible serine protease PMSF and EDTA suggests the degradation may be caused by a combination of a serine and metal-dependent protease.

While evidence points to a either a serine or metal-dependent protease, the species mediating the observed degradation remains unclear. Prior to observing ParD for degradation, extensive purification was performed. ParD was initially purified by IMAC chromatography using a Ni-NTA column; the protein was then desalted to remove imidazole and was treated with thrombin for removal of the N-terminal 6x-His tag. After thrombin cleavage, ParD was passed again over a Ni-NTA column and collected in the flow-through, with any residual contaminants from the first Ni-NTA purification expected to also be retained in this subsequent Ni-NTA purification step. A final size-exclusion step was included to separate ParD from thrombin. Despite these extensive purification steps, purification alone was not sufficient in blocking ParD degradation, and analysis by direct-infusion mass spectrometry identified peaks attributed to only ParD. The mass spectrometry results reveal that the toxin-binding region is completely absent from degraded antitoxin, and the DNA binding region remains intact. The mass spectrometry results combined with our *in vitro* degradation assays suggest *Pseudomonas aeruginosa* ParD may have latent protease activity that is capable of autocatalytic self-cleavage. If this is also the case *in vivo*, it indicates that it may be possible for the cell to uncouple the transcriptional repression functionality of ParD from its toxin neutralizing function. While this analysis did not provide answers to what was mediating the degradation, as no peaks of significant height were detected other than antitoxin, it does bring an interesting idea of uncoupling toxin binding with DNA binding in an antitoxin molecule, and suggests a potential protease activity for at least some ParD antitoxins.

## Experimental procedures

### Expression and purification of *Pseudomonas aeruginosa* ParD and ParE

Purified ParD and ParE protein were obtained by following a previously published protocol (19). To obtain ParDE complex, ParD and ParE were buffered exchanged into 50 mM MES pH 6.0, 300 mM NaCl, 5% glycerol, then mixed in equimolar ratios. The mixed proteins were incubated on ice for 30 minutes prior to purification by size-exclusion chromatography using a Superdex 75 10/300 GL column (GE Healthcare) pre-equilibrated with a 50 mM MES pH 6.0, 300 mM NaCl, 5% glycerol buffer.

### *In vitro* degradation Assay

Purified samples at 300 μM in 50 mM Tris pH 8.5, 300 mM NaCl buffer were incubated as indicated; at designated time points 25 μL was removed, combined with reducing SDS loading dye, and heated at 95°C for 5 minutes. For samples treated with PMSF (1 mM), or protease inhibitor cocktail (Sigma, used at a final 1x concentration), the compounds were added during lysis. EDTA was added to samples after the initial Ni-NTA purification step. PMSF and EDTA were also maintained in purification buffers, while additional protease inhibitor cocktail was added after the final size-exclusion step. Once all time points for a series were collected, samples were loaded onto 12% acrylamide gels with a Tris-Tricine buffer system (20), separated by electrophoresis, and visualized by staining with Coomassie blue. Relative band intensities were measured using ImageJ (21) (see **Figs 1-3**).

### Direct Infusion Mass Spectrometry

Mass spectrometry of intact and degraded WT ParD was performed with a Orbitrap Fusion Tribrid Mass Spectrometer (Thermo Fisher Scientific). Prior to analysis by mass spectrometry, samples were desalted into a 50mM Ammonium bicarbonate pH 6.0 buffer and concentrated to 1 mg/ml. Samples were mixed 1:1 with a solution of 30% methanol, 0.1% acetic acid prior to direct injection into the electrospray ionization source. Data collection was performed using both the ion trap and Orbitrap mass analyzers. Mass spectra were analyzed using the Xcalibur software package (Thermo Fisher Scientific) (see **Figs 5-6**).

## Acknowledgements

We appreciate the assistance of the OU Protein Production and Characterization core facility for assistance with protein purification. The OU Protein Production and Characterization Core facility is supported by an Institutional Development Award (IDeA) from the National Institute of General Medical Sciences of the National Institutes of Health under grant number P20GM103640. We appreciate the expert assistance of Dr. Steve Hartson for mass spectrometry data collection and analysis, which was carried out at the OSU DNA/Protein Core Facility at Oklahoma State University, Stillwater OK.

## Author Contributions

KJS collected all data; both authors participated in the project design, data analysis, manuscript drafting and editing, and have approved the submitted version. Funding Research reported in this publication was supported by an Institutional Development Award (IDeA) from the National Institute of General Medical Sciences of the National Institutes of Health under grant number P20GM103640 (PI: A West, Project Leader: CRB), and an award for project number HR17-099 from the Oklahoma Center for the Advancement of Science and Technology (to CRB). The content is solely the responsibility of the authors and does not necessarily represent the official views of the National Institutes of Health.

